# Trade-offs, trade-ups, and high mutational parallelism underlie microbial adaptation to extreme feast/famine

**DOI:** 10.1101/2023.10.04.560893

**Authors:** Megan G. Behringer, Wei-Chin Ho, Samuel F Miller, Sarah B. Worthan, Zeer Cen, Ryan Stikeleather, Michael Lynch

## Abstract

Microbes are robust organisms capable of rapidly adapting to complex stress, enabling the colonization of harsh environments. In nature, microbes are regularly challenged by starvation, which is a particularly complex stress because resource limitation often co-occurs with changes in pH, osmolarity, and toxin accumulation created by metabolic waste. Often overlooked are the additional complications introduced by eventual resource replenishment as successful microbes must withstand rapid environmental shifts before swiftly capitalizing on replenished resources to avoid invasion by competing species. To understand how microbes navigate trade-offs between growth and survival, ultimately adapting to thrive in environments with extreme fluctuations, we experimentally evolved 16 *Escherichia coli* populations for 900 days to repeated feast/famine cycles of 100-day starvation before resource replenishment. Using longitudinal population-genomic analysis, we found that evolution in response to extreme feast/famine is characterized by narrow adaptive trajectories with high mutational parallelism and notable mutational order. Genetic reconstructions reveal that early mutations result in trade-offs for biofilm and motility but trade-ups for growth and survival, as these mutations conferred correlated advantages during both short-term and long-term culture. Our results demonstrate how microbes can navigate the adaptive landscapes of regularly fluctuating conditions and ultimately follow mutational trajectories that confer benefits across diverse environments.

## Introduction

In natural environments, the availability of resources is often inconsistent, posing a constant adaptive challenge for microbial populations across diverse ecosystems, including the soil, aquatic habitats, and the human gut [1–3]. These periodic fluctuations in resource availability, also referred to as feast/famine, demand microbial populations to survive varying durations of starvation but also to swiftly adjust and efficiently utilize resources upon replenishment to compete with neighboring species. Host-associated microbes and facultative pathogens are under particularly pronounced pressure, as these organisms must navigate feast and famine conditions across wide-ranging timescales to support their multi-environment life histories [4–8].

Investigation of microbial adaptations to starvation conditions has revealed significant tradeoffs between growth and survival [9–12]. In studies of starvation without resource replenishment, a typical strategy involves evolving alterations to metabolism that relieve catabolite repression, allowing microbes to survive on second-tier carbon sources and resources released from dead cells [13, 14]. However, as starvation continues and conditions intensify, populations can become over-adapted to starvation conditions and maladapted to resource replenishment conditions. Starvation-induced mutagenesis may further increase the potential for maladaptation by contributing to deleterious mutation loads [15–19]. Thus, strategies that reduce maladaptation to resource replenishment conditions should increase in importance as the duration of famine continues [20–24]. One extreme example of a survival strategy for long-term starvation is endospore formation in Firmicutes, where species like *Bacillus subtilis* can develop into a dormant cell type, almost completely inactivating metabolism, to resist very harsh environments [25]. Yet, endospore formation is not immune to trade-offs [26, 27], as prolonged germination times can inhibit cells from rapidly capitalizing on resources. Firmicutes that have lost the ability to form endospores instead prioritize metabolite transport, resulting in decreased ability to withstand stress but an increased capability to colonize new environments [28].

In contrast to the situation in sporulating bacteria, the possible adaptive trajectories for non-sporulating bacteria to navigate trade-offs while adapting to feast/famine fluctuations are less clear [29–32]. For example, *Escherichia coli* can colonize a broad habitat range, including soil, wastewater, and the lower gut. Prior studies have hypothesized that *E. coli*’s ability to scavenge carbon from a wide variety of nutrient resources [33] may contribute to extended survival in low-nutrient environments, as reports have documented *E. coli* to survive over 260 days in freshwater and over nine years in soil [34, 35]. Despite *E. coli*’s varied life history, most investigations have focused solely on adaptation to prolonged starvation [36–39] or feast/famine cycles with relatively brief durations of famine [40–42]. Empirical investigations of *E. coli* have revealed that populations surviving periods of prolonged starvation often accumulate mutations in the genes *rpoS* [43] and *lrp* [44], which contribute to a phenotype known as “growth advantage in stationary phase” (GASP) [45, 46, 37]. As resource limitation becomes more severe, mutations in subunits of the RNA polymerase holoenzyme (*rpoB* and *rpoC*) confer further benefits, as do mutations in the enzyme that catalyzes the hydrolysis of cyclic AMP (*cpdA*), a resource-availability signaling molecule [38, 47]. However, an understanding of how these genetic and phenotypic responses generalize or scale across different durations of resource limitation is still lacking, and it is unknown if these evolved genotypes and phenotypes will still be observed when an influx of replenished resources occurs periodically. Thus, studies of microbial evolution in the context of fluctuating feast/famine cycles with various magnitudes and periodicities are needed.

To address these issues, we surveyed the evolutionary response of *E. coli* populations under 100-day feast/famine cycles, where the evolving populations are transferred into fresh lysogeny broth (LB) every 100 days. Evolving populations were established from ancestral lines with two different initial genetic backgrounds: a wild-type strain (WT) and a WT-derived strain with impaired methyl-directed mismatch repair (MMR-) wherein an engineered deletion of *mutL* yields a ∼100× increase in the mutation rate [48, 49]. Each genetic background was replicated in eight parallel experiments, resulting in 16 (8×2) experimental populations. Longitudinal metagenomic sequencing of evolving populations over 900 days allowed us to investigate how the initial mutation rate and spectrum affect evolution, and if repeated famine results in genetic and phenotypic parallelism between initial genotypes. We can then compare the genotypic parallelism observed in response to 100-day feast/famine cycles to those in the prior investigation involving much shorter feast/famine cycles [50]. Lastly, genomic analysis with genetic reconstruction and phenotyping allowed us to probe the biological relevance and tradeoffs associated with early adaptive mutations.

## Results

### Ecotypic diversification is uncommon in 100-day feast/famine cycles

Eight WT and MMR-populations of *Escherichia coli* were experimentally evolved in 16 × 100 mm glass culture tubes enduring nine feast/famine cycles, each consisting of 100 days of starvation before transferring 1 mL of culture into 10 mL fresh LB broth. The genomic evolution of these experimental populations was tracked with metagenomic sequencing on 1 mL samples of fully vortexed pre-transfer cultures every 100 days (mean coverage > 100×, **Table S1)**. In total, we identified 16,886 mutations and observed linear accumulation of new mutations throughout the evolution experiment with 4,963 mutations remaining at Day 900. As expected, the rate of genome evolution is faster for MMR-populations than WT populations (**Fig. 1A**), consistent with the earlier observation that the MMR-populations decreased their mutation rates by ∼50% under the evolution in 100-day feast/famine cycles but still retained ∼21× higher mutation rates than the evolved WT populations [49]. Because we have previously observed the evolution of ecotypes in a heterogeneous culture environment with shorter durations between transfers into fresh media (1-day feast/famine[42] or 10-day feast/famine[50]), we looked to see how the evolutionary response to 100-day feast/famine cycles affected ecotypic diversification. We applied a clade-aware hidden Markov chain (caHMM) to our metagenomic sequencing data to identify evidence suggesting the presence of coexisting clades and estimate the maximum duration of clade coexistence [51]. Only 2 of 16 evolving populations produced evidence of multiple divergent clades coexisting longer than three 100-day feast/famine cycles (**Fig. 1B**). This suggests that instead of ecotypic diversification, successive selective sweeps characterize early adaptation to 100-day feast/famine cycles and that strong selection is shaping the evolving populations.

**Figure 1.**
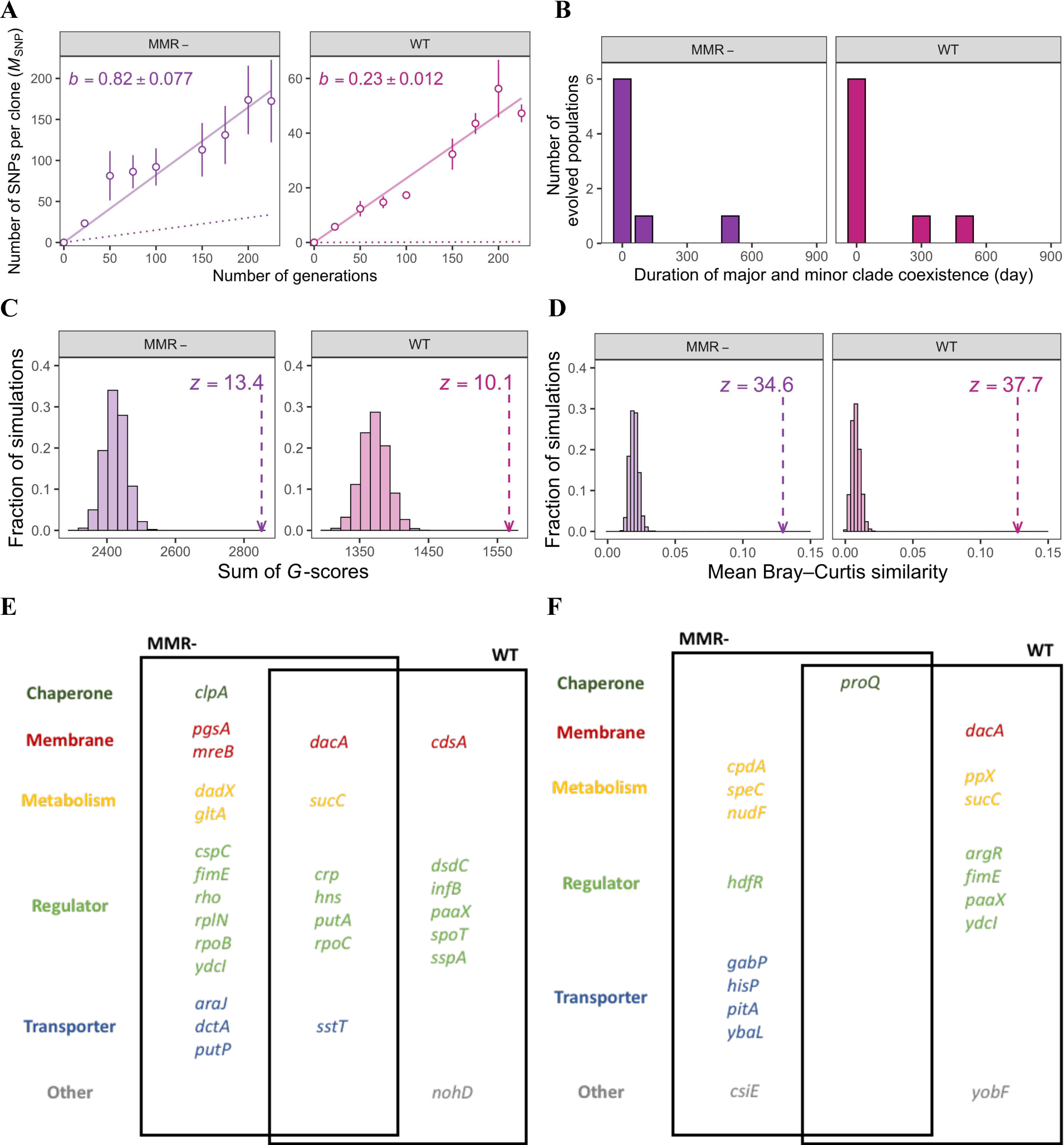
Genomic evolution in the populations with either methyl-directed mismatch-repair defected (MMR-) or wild-type (WT) background shows parallelism of fixed mutations. **(A)** The single-nucleotide polymorphism (SNP) accumulated through time (in generations) in either genetic background. Each dot represents the average number of SNPs found in a cell for an evolved population; the accompanied error bar represents ± s.e.m.. The solid line shows the linear-regression result of the mean number of SNPs against the generation numbers; the slope (*b*) and the associated S.E. are also printed. The dashed line shows the predicted numbers of SNPs based on the initial mutation rates. **(B)** No strong evidence for subpopulational structures in the evolved populations, as the hidden Markov model estimated a very short duration for the coexistence of different clades in an evolved population. **(C)** The parallelism of fixed nonsynonymous mutations among the evolved populations in either genetic background is statistically significant based on the measurement of the sum of *G*-scores. The arrow represents the observed value from the evolved populations, whereas the histogram shows the null expectation with no parallelism. The significance of the observed value is evaluated by comparison to the mean and normalization by the standard deviation of the null expectation, resulting in the noted *z* score. **(D)** The parallelism of fixed nonsynonymous mutations among the evolved populations in either genetic background is statistically significant based on the mean Bray-Curtis similarity among all the population pairs. The arrow represents the observed value from the evolved populations, whereas the histogram shows the null expectation with no parallelism. The significance of the observed value is evaluated by comparison to the mean and normalization by the standard deviation of the null expectation, resulting in the noted *z* score. **(E)** This list of genes that are significantly enriched by the fixed nonsynonymous mutations in the evolved populations. **(F)** This list of genes that are significantly enriched by the fixed structural or nonsense mutations in the evolved populations.

### Prolonged famine results in significant parallelism in fixed mutations

The mode of natural selection and the availability of adaptive trajectories can be described by investigating mutational parallelism, i.e., the tendency for mutations to repeatedly fix in the same gene across replicate populations [52]. To quantify mutational parallelism across evolving populations and identify gene candidates in which acquired mutations likely represent molecular adaptations to 100-day feast/famine cycles, we employed two metrics to rank selective targets based on an overrepresentation of fixed nonsynonymous mutations: sum of *G*-scores [53] and mean Bray–Curtis similarity [54]. We initially focused on fixed nonsynonymous mutations, as these have potential to alter protein functions in evolved populations. If fixed non-synonymous mutations are concentrated in a smaller subset of genes, the resulting values for these two metrics will be higher. For both MMR- and WT populations, both the sum of *G*-scores and the Bray-Curtis similarity values were significantly greater than expected based on a simulated null distribution (**Fig. 1C-D**), revealing a significant amount of evolutionary convergence. The adaptive trajectories for 100-day feast/famine cycles are exceptionally similar, as the level of statistical significance (*z* score) for the observed parallelism for 100-day feast/famine cycles is much higher than those for 1-day feast/famine cycles [50].

Identifying mutational parallelism also allows us to distinguish which mutations are likely to be under positive selection and which are likely being promoted to fixation by genetic draft [55], as individual genes enriched for nonsynonymous mutations will have greater individual *G*-scores. Across both MMR- and WT populations, we identified 28 genes as enriched for nonsynonymous mutations (**Fig. 1E**; **Table S2**). Because disruptive ‘structural’ mutations (such as indels, IS-element insertions, or nonsense mutations) can also greatly impact a gene’s function, we additionally looked for parallelism in fixed structural mutations and identified 18 genes using the same method (**Fig. 1F; Table S3**). In total, we identified mutational parallelism for 41 genes across the nonsynonymous and structural mutation gene sets, with five genes (*dacA*, *fimE*, *paaX*, *sucC*, and *ydcI*) occurring in both gene sets. Given the frequency and breadth of mutations in these five genes, total loss, or severe reduction in the function of these genes is likely to be highly adaptive for 100-day feast/famine cycles. Two of these five genes (*fimE* and *ydcI*) were also candidates for the adaptive to 1-day feast/famine cycles [50], suggesting that they may provide benefits during both starvation and resource replenishment conditions (**Fig. S1**).

### Regulator, transport, and cell wall/membrane genes are enriched for parallel variants

Examination of annotated gene functions and targeted pathways or processes revealed that nonsynonymous and structural mutations in the experimental lines commonly affect genes associated with five major classes: chaperones (2 genes), cell wall/membrane maintenance (4 genes), regulation of global gene expression (17 genes), metabolism (7 genes), and ion-driven transport (8 genes) (**Fig. 1 E-F**). Across these genes we observe significant mutational order, or high parallelism in at which time point a mutation is first detected within an individual gene during evolution (ANOVA, F = 4.32, P = 2.35 x 10^−12^; **Fig. 2A**). This signal of parallelism is present across WT and MMR-backgrounds (ANOVA, WT: F = 2.33, P = 1.31 x 10^−3^; MMR: F = 3.29, P = 4.76 x 10^−7^; **Fig. S2**) and is most significant for genes associated with regulation, although transport-associated genes also exhibit significant mutational order (ANOVA, regulators: F = 5.87, P = 3.21 x 10^−9^; transporters: F = 2.76, P = 0.01; **Fig. S3**).

**Figure 2.**
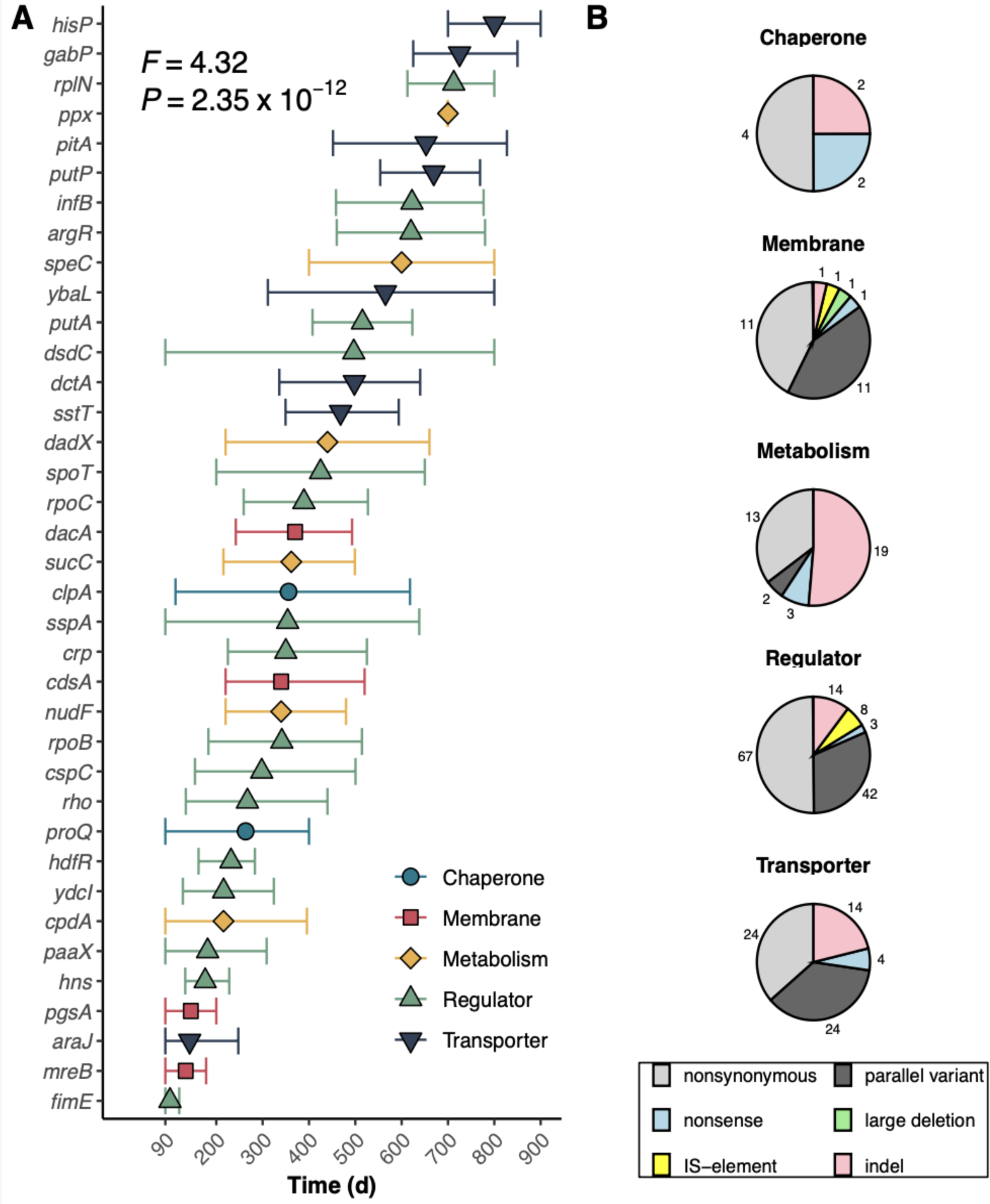
Timeline and distribution of mutations in significantly enriched genes. A) Timeline illustrating the timepoint when a mutation is first observed in a significantly enriched gene. For each population, only the first fixed mutation was considered. Individual data points represent the mean timepoint a mutation is first detected, and error bars represent the 95% CI of the mean. Colors and shapes indicate the assigned functional group of each gene. B) Pie charts illustrating the relative distribution of mutation types detected in significantly enriched genes. Colors indicate the mutation type and numbers indicate the number of each mutation type observed.

Within regulation-associated genes, we can further define mutational order that appears to be based on downstream gene functions. Genes annotated as associated with the decision between committing to biofilm formation and motility (e.g., *fimE*, *hns*, *hdfR*) are the earliest mutations to be promoted by selection, typically within the first 300 days. Mutations that directly affect transcription (*rho*, *rpoB*, *rpoC*) follow in the second 300 days, while mutations directly affecting translation (*infB*, *rplN*) ultimately arise in the last 300 days of experimental evolution. We also noted that different mutation types tend to fix in regulation, transport, and membrane-associated genes in particular orders. Nonsynonymous SNPs tend to fix later than nonsense SNPs and IS-element insertions in regulator genes (ANOVA, F = 2.67, P = 0.050; **Fig. S3B**) and may be fine-tuning regulator activity after early loss-of-function mutations of large effect. However, in contrast to the situation with regulators, nonsynonymous SNPs occur earlier in transport genes than structural mutations, and there is no significant order observed in the timing of selection on membrane-associated genes based on mutation type (ANOVA, transporter: F = 4.97, P = 3.72 x 10^−3^; membrane: F=0.722, P = 0.61; **Fig. S3D, F**). Transport-associated genes also have a higher enrichment of loss-of-function mutations than regulators, with the highest proportion of loss-of-function mutations observed in genes associated with metabolism (metabolism: 59%; transport: 27%; regulators: 18%, membrane: 15% of mutations), which may mitigate energetic-waste losses by limiting the expression of unnecessary genes during starvation conditions.

Another aspect of parallelism that defines adaptation to 100-day feast/famine cycles is the high proportion of parallel variants observed in regulation, transport, and membrane-associated genes (**Table S4**; **Fig. 2B**). We define parallel variants as identical nonsynonymous SNPs that independently occur at the same nucleotide position in two or more replicate populations. Parallel variants occurred in 12 of the 17 regulation-associated genes enriched for nonsynonymous mutations, accounting for 38.5% (42 out of 109) of the nonsynonymous SNPs and 31.3% of all mutations observed in these genes after 900 days. In transport- and membrane-associated genes, parallel mutations accounted for 50% of the nonsynonymous SNPs (transporters: 24 out of 48 SNPs, 4 out of 8 genes; membrane: 11 out of 22 SNPs, 4 out of 4 genes). Because parallel variants commonly arose in functional regions of genes, we expect their fixation to be a product of strong selection for altered protein function.

### Early mutations result in tradeoffs between biofilm and motility without costs to growth or survival

As early mutations in regulation-associated genes often resulted in the loss of function of genes associated with the decision between committing to biofilm formation and motility, we measured biofilm and motility activity for evolved populations to identify any phenotypic patterns [56, 57]. Overall, WT and MMR-populations both evolved significant increases in biofilm formation during 100-day feast/famine cycles (Pairwise t-test, WT: P = 1.28 x 10^−8^; MMR-: P = 9.70 x 10^−2^) and MMR-populations exhibited significant reductions in motility (Pairwise t-test, WT: P = 0.11; MMR-: P = 0.023; **Fig. 3 A-B, Fig. S4**). Thus, to determine how each of the early loss of function mutations in regulator genes contribute to the observed biofilm and motility phenotypes, we deleted from the experimental ancestor each of the regulator genes that commonly evolved a loss of function mutation within the first 300 days (*fimE*, *hdfR*, *paaX*, and *ydcI*) and repeated the biofilm and motility assays. Consistent with the phenotypic pattern observed in evolved populations, the deletion of *fimE* causes a significant increase in biofilm formation and a decrease in motility (ANOVA with Tukey’s HSD, *fimE_Biofilm_*: P = 0.001; *fimE_Motility_*: P = 0.061). Deletions of *hdfR*, *paaX*, or *ydcI* did not induce biofilm formation but significantly increased motility (ANOVA with Tukey’s HSD, *hdfR_Motility_*: P = 0.022; *paaX_Motility_* P = 9.72 x 10^−4^; *ydcI_Motility_* P = 0.015; **Fig. 3C-D**).

**Figure 3.**
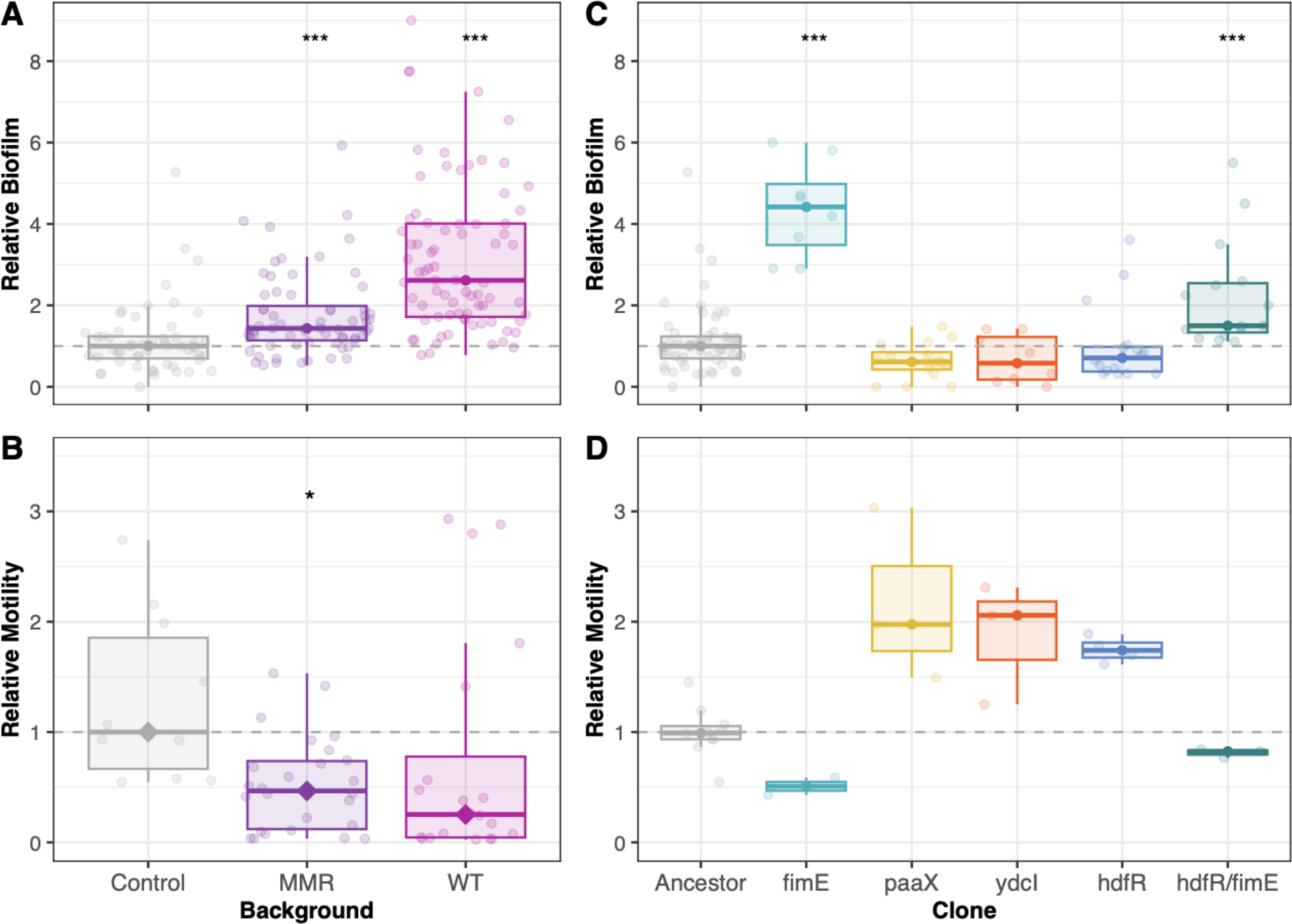
Relative biofilm and motility activity produced by evolved populations and engineered deletion strains. After 900 days of evolution, populations exhibited changes in A) biofilm production (measured by absorbance at 550 nm) and B) motility (measured by culture radius). Deletion of early regulator genes demonstrates the contribution of loss of function mutations to C) biofilm and D) motility phenotypes. Individual data points denote the relative deviation of the measured phenotype for each replicate from the median ancestral value. Boxplots illustrate the quartile ranges of the data. Colors differentiate the genetic backgrounds of the evolved populations or deletion strains. Significance of pairwise comparisons with the ancestor are denoted by asterisks (*: P < 0.05; **: P < 0.01; ***: P < 0.001).

Biofilm formation and motility commonly exhibit trade-offs, as motility can destabilize biofilms and result in dispersal [58, 59, 12, 60], and the molecular mechanisms regulating these two traits are often associated [61]. Comparing our measurements of the relative changes in biofilm formation and motility for the *fimE*, *hdfR*, *paaX*, and *ydcI* deletion mutants confirms a trade-off between biofilm and motility (Pearson’s *r* = −0.891, P = 0.042; **Fig. 4A**). As prior studies also identified growth and survival as traits that exhibit trade-offs [62, 10, 63–65], we considered how these early loss-of-function mutations might contribute to fitness throughout a cycle of growth and starvation. We conducted co-culture competition assays between the deletion mutants and the WT ancestor, accessed their frequency after 1, 4, and 10 days of famine, and calculated their fitness in terms of selection rate (**Fig. 4B-D**). Competitions revealed that all deletion clones, except for Δ*paaX,* demonstrated positive selection rates at all time points resulting in a positive correlation between growth after 1 day of culture and survival over 10 days of culture (Pearson’s *r* = 0.922, P = 0.026; **Fig. 4B**), illustrating that growth/survival tradeoffs can be navigated by single loss-of-function mutations without a negative effect on either trait.

**Figure 4.**
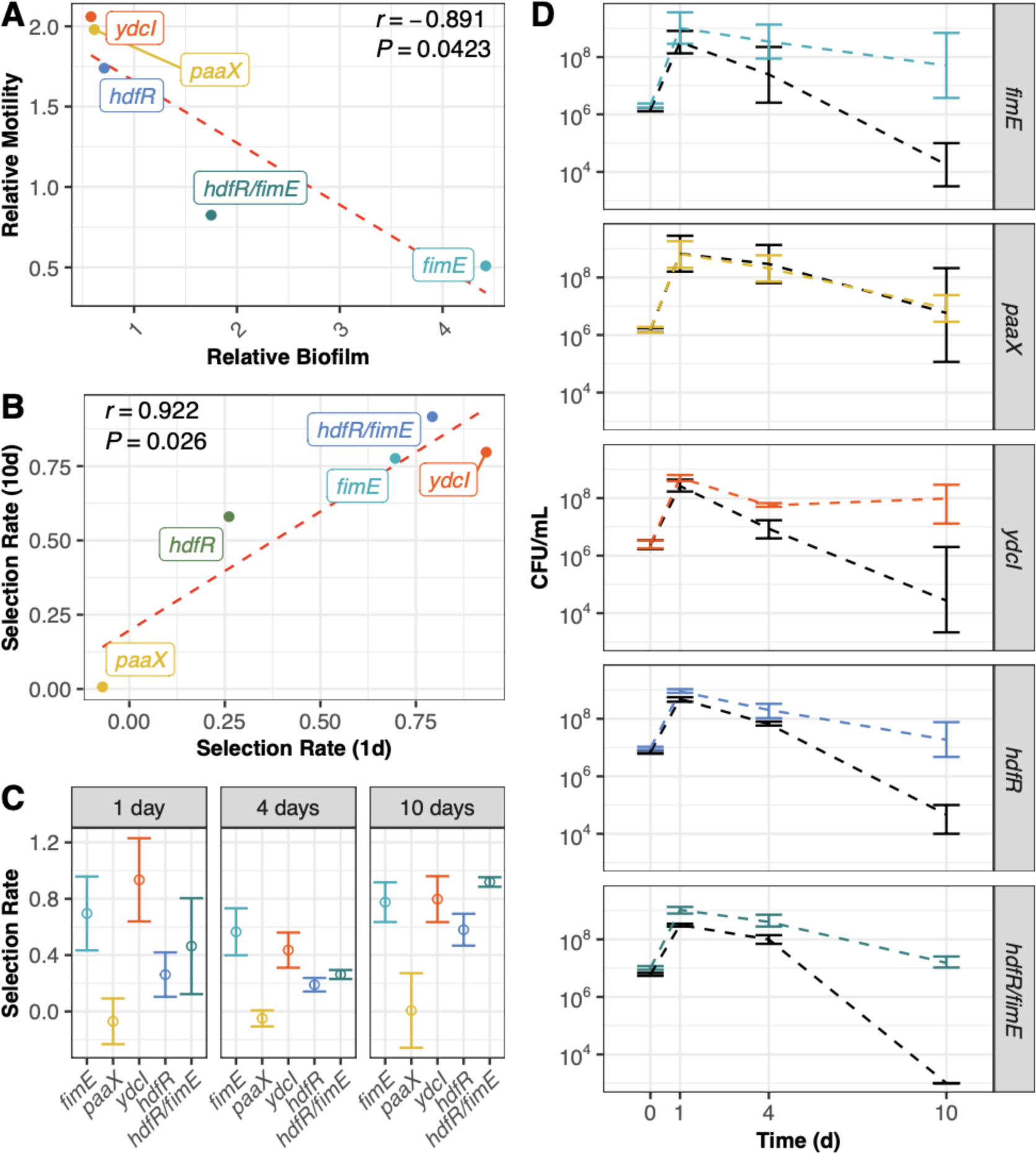
Early loss of function mutations in regulation-associated genes exhibit trade-offs in biofilm/motility but not growth/survival. Scatter plots illustrate that A) median relative motility and median normalized biofilm phenotypes of engineered deletion strains are negatively correlated, but B) selection rate during growth (1-day culture) and selection rate during survival (10-day culture) for engineered deletion strains is positively correlated. For panels A and B, the dashed red line represents the linear fit of the data. C) Engineered deletion strains exhibit positive selection rates across 1-, 4-, and 10-days of culture without resource depletion when competed against WT ancestor suggesting that early loss of function mutations confer generally positive effects on growth and survival. Open circles denote mean selection rates (differences of Malthusian parameters before and after the experimental evolution) and error bars represent the 95% CI of the mean. D) CFU counts of engineered deletion strains and the WT ancestor when cocultured for 1-, 4-, and 10-days without resource repletion. Lines connect the mean CFU counts for each competitor at 1-, 4- and 10-days, and error bars represent the 95% CI of the mean. For all panels, colors denote the different engineered deletion strains while black denotes the WT ancestor.

Despite the conflicting biofilm/motility phenotypes generated by early loss-of-function mutations in regulation-associated genes, these mutations regularly co-occur in 100-day feast/famine populations, as 11 of the 16 populations containing a fixed biofilm-enhancing mutation in *fimE*, also have at least one fixed motility-promoting mutation in *hdfR*, *paaX*, or *ydcI* (Hypergeometric test, *P* = 0.00023). To further investigate the phenotypic effects of these opposing mutations, we engineered a double mutant combining a motility-promoting mutation (Δ*hdfR*) and a biofilm-enhancing mutation (Δ*fimE*). We focused on *hdfR* and *fimE* because six populations evolved fixed mutations in these genes by the 300-day time point. Biofilm and motility assays for the Δ*hdfR*/Δ*fimE* double mutant confirmed the trade-offs predicted from the phenotypes of the single mutants as the double mutant exhibits additive effects, producing less motility but more biofilm than the Δ*hdfR* mutant (ANOVA with Tukey’s HSD, Motility: *P* = 0.027; Biofilm: *P* = 0.023) (**Fig. 3C-D, Fig. 4A**). Next, we wanted to confirm that the Δ*hdfR*/Δ*fimE* double mutant is still able to navigate the growth/survival tradeoff without negatively affecting either trait as observed in the Δ*hdfR* and Δ*fimE* single mutants. Co-culture of the Δ*hdfR*/Δ*fimE* double mutant with the WT strain revealed that the double mutant exhibits similar fitness as the single mutants during 1-day growth and the greatest fitness of all the engineered deletion mutants after 10 days of starvation. Thus, while the Δ*hdfR*/Δ*fimE* double mutant exhibits intermediate biofilm and motility phenotypes, the strict trade-offs between biofilm and motility do not appear to extend to fitness across feast/famine conditions. Lastly, it is possible that although deletion of *hdfR*, *paaX*, or *ydcI* confers increased motility, motility may not be the primary phenotypic trait that is under selection as these regulators may have broader effects. Further investigation of these regulators is needed to resolve the phenotypes produced in different genetic contexts and how mutations in transcriptional regulators contribute to adaptation during long-term feast/famine cycles.

## Discussion

This study reports on the evolution of *E. coli* populations cultivated in repeated 100-day feast/famine cycles over 900 days. Compared to results of experimental evolution with daily resource replenishment, populations evolving in the more-extreme 100-day feast/famine cycles exhibited little to no ecotypic diversification. Strikingly, we observed significant parallelism in fixed mutations during adaptation to repeated extreme long-term starvation, suggesting that strong selection shapes these populations. Superficially, identifying gene-level parallelism in fixed mutations is not surprising; one of the common goals of adaptive laboratory evolution experiments is to assess the repeatability of evolutionary processes [66]. Gene-level parallelism is common in microbial populations evolving to standard laboratory conditions [53, 67] and more stressful conditions such as stress due to temperature [68], antimicrobials [69], starvation [41, 50], and reactive oxygen [70]. However, a combination of gene-level parallelism with significant mutational order is a unique characteristic defining evolution to extreme feast/famine cycles. Mutation order [71] has contributed to several evolutionary processes including speciation [72], tumorigenesis [73], and adaptation to stress [74], and can provide information about the adaptive landscape [75]. In the lab, *S. cerevisiae* and *E. coli* populations exhibit mutational order during experimental evolution to antimicrobial stress [76, 77], but this pattern is restricted to the few very early first-step mutations needed to confer antimicrobial resistance.

Through longitudinal sequencing of *E. coli* populations evolving to 100-day feast/famine cycles, we identified strong and convergent patterns of mutational order that extend throughout the 900-day experiment and are particularly strong among mutations affecting global regulators. Here, mutations appear to arise in three broad batches beginning with regulators that control biofilm and motility within the first 300 days, followed by core transcriptional genes, and mutations affecting core translational genes arising last. The order of these broad categories may indicate the processes whose alteration provides the greatest environmental-dependent effects on fitness. A prior study of mutations affecting *E. coli* growth behavior during batch culture demonstrated that in less preferred carbon sources, like succinate, mutations that shorten lag phase tend to sweep to high frequencies before mutations that increase the maximum growth rate [78]. As lag phase during growth on succinate typically lasts 15 h, mutations that can shorten lag phase to 5 h provide greater benefits than what would be achieved by modestly increasing growth rates. However, in lactose, where the initial duration of lag phase is much shorter, the order of mutations switches, and increased growth rates are prioritized instead. Thus, we expect a similar framework during long-term starvation in culture tubes. For instance, if access to oxygen and migration to the surface-air interface provides the greatest early fitness benefits, then mutations influencing biofilm formation should arise first. Despite starvation commonly being thought of as a signal that triggers the dispersal of biofilms, biofilm formation can also be an adaptive strategy for microorganisms that increases survivability in starving conditions [79]. We observe that loss of function mutations in a negative regulator of biofilm-formation, *fimE,* typically arise within the first 100-days of experimental evolution to extremely long feast/famine cycles (**Fig. 2**). By deleting *fimE* from the experimental ancestor, we confirmed that these mutations not only confer increased biofilm production but also increased fitness in both the growth and survival phases of feast/famine cycles. Thus, increased biofilm formation is likely a critical early step for adaptation to long-term feast/famine cycles.

Another documented response to starvation is the expression of flagellar motility genes by *E. coli* during the early nitrogen starvation response [80]. However, we were surprised to observe the quick fixation of loss-of-function mutations that increase motility (Δ*hdfR,* Δ*ydcI,* and Δ*paaX*) occurring alongside the loss-of-function mutations that increase biofilm formation (Δ*fimE*), as these two processes are typically counter-regulated. Disruption of the *hdfR* and *ydcI* genes in the experimental ancestor confers similar fitness increases as the Δ*fimE* mutant, and mutations in *hdfR* and *ydcI* always arise in tandem with mutations in *fimE* during experimental evolution. As a genotype’s fitness manifests through its associated phenotype, it is interesting to observe that the Δ*hdfR*/Δ*fimE* double mutant exhibits muted biofilm/motility phenotypes but greater fitness values than the Δ*hdfR* and Δ*fimE* single mutants. One possibility is that loss of function mutations in *hdfR* are compensatory and help recover some fraction of the ancestral motility phenotype lost due to mutation in *fimE.* However, this does not entirely explain why strains without *fimE* mutations exhibit similarly diminished motility. Another possibility is that the motility phenotype produced by loss of function mutations in *hdfR* is incidental due to pleiotropy, and motility is not the actual phenotype under selection. This explanation is consistent with the motility phenotypes produced after the deletion of *paaX* and *ydcI* - two genes that are not typically associated with motility.

There are two other possibilities for why loss-of-function mutations in *hdfR* would be beneficial during starvation conditions. The first of which is glutamate synthesis. In *E. coli*, glutamate metabolism is highly important, as up to 80% of *E. coli*’s nitrogen flows through glutamate [81] created through either the ammonia assimilation cycle III pathway via glutamate synthase (GltBD) or through the L-glutamate biosynthesis I pathway via glutamate dehydrogenase (GdhA). HdfR upregulates GltBD, generating L-glutamate during nitrogen limitation through an ATP-dependent process that recycles ammonia. In the absence of GltBD and when ammonia is abundant, *E. coli* uses GdhA to generate L-glutamate by incorporating ammonia in an ATP-independent process. Because *E. coli* cultures in LB broth typically cause alkalization of the media due to excreted ammonia, disruption of *hdfR* may encourage glutamate synthesis through the L-glutamate biosynthesis I pathway, thus conserving ATP and limiting ammonia accumulation.

Previously, we reported that 1000 days of evolution in response to 100-day feast/famine cycles conferred an average mutation rate reduction of 50% for MMR-populations [49]. As such, a second possible explanation for why we observe loss-of-function mutations in *hdfR* could be due to a selective pressure to decrease mutation rates. Prior studies report Δ*hdfR* mutants to confer a 6 to 9-fold reduction in mutation rate [82] and overexpression of *paaX* to confer greater than 100-fold increases in mutation rate [83], suggesting that mutations in these genes may have broad physiological effects. Comparing populations based on the number of SNPs that reach a frequency of greater than 20% indicates that MMR-populations containing mutations in *paaX* or *hdfR* evolve fewer SNPs during 900 days of experimental evolution (avg. 40% reduction, *P* = 0.006, t-test; **Fig. S5**). However, this pattern is not significant for WT backgrounds where the base genetic mutation rate is considerably less (*P* = 0.15, t-test; **Fig. S5**). Reduction of mutation rates, especially during the extended famine portion of the feast/famine cycle, may help regulate how new genetic variation is introduced and subsequently filtered by natural selection throughout the culture cycle. Otherwise, mutations induced by starvation-induced mutagenesis [84] might greatly outweigh replication-induced mutations, potentially resulting in the population becoming more adapted to the famine phase at the expense of adaptation to the feast and resource repletion phase. A potential second-order effect on mutation rates may also explain why the single Δ*paaX* mutation failed to confer any fitness effects despite loss-of-function mutations in *paaX* fixing in 7 out of the 16 populations. Alternatively, *paaX* mutations may confer benefits that are dependent on an earlier arising mutation or adaptive for the more extreme starvation conditions that arise beyond the 10 days assessed in competition assays.

Lastly, preventing maladaptation to the resource-replenishment conditions may also explain why adaptation to 100-day feast/famine cycles does not result in the fixation of mutations in the global regulator genes that are more commonly discussed in the context of long-term starvation without large-scale resource repletion, like *rpoS* [36, 43]. Attenuated *rpoS* activity can contribute to increased survival in starvation conditions through increased tolerance of alkaline pH and by relaxing catabolic repression allowing *E. coli* to simultaneously co-utilize amino acids and second-tier carbon sources. However, relaxing catabolic repression may have consequences during resource replenishment, as cells will have a reduced ability to prioritize preferred carbon sources and would be at risk of being outcompeted. As such, the genotypes and phenotypes that are adaptive in environments with regular feast/famine cycles have the potential to vary greatly based on the duration of starvation and the magnitude of resource replenishment. A comparison of genes overrepresented for mutation in this study with a study of 10-day feast famine cycles finds eight gene targets (*argR*, *crp*, *dacA*, *pitA*, *putA*, *putP*, *rpoB*, and *sstT*) in common [50]. Thus, further investigation examining adaptation to starvation across conditions, timescales, and fluctuation frequency would contribute to a more universal understanding of the adaptation to resource limitation and the tradeoffs involved.

## Materials and Methods

### Construction of ancestral strains and experimental evolution

Experimentally evolved populations were propagated from the following ancestor strains: PFM2, a prototrophic derivative of *E. coli* K-12 str. MG1655 (denoted as WT) and PFM5, which is PFM2 with gene deletion of *mutL* (denoted as MMR-). To identify any cross-contamination between evolving populations during the evolution experiment, a neutral marker consisting of a 3513-bp deletion in the *araBAD* operon (*ara-*) was introduced in both ancestor strains and used in half of the experimental populations (4 out of 8 WT and 4 out of 8 MMR-). To screen for contamination, we streaked experimental populations every 100 days on MacConkey agar (BD Difco) supplemented with 0.4% arabinose.

Evolution experiments were initiated by inoculating a single-isolated progenitor colony cultivated overnight at 37 °C on LB agar plates in 10 mL of LB-Miller broth (Dot Scientific). We then maintained the cultures in 16-× 100 mm glass tubes shaking at 175 rpm at 37°C. Every 100 days, we thoroughly vortexed the culture and transferred 1 mL of the vortexed culture into 10 mL fresh LB broth. Copies of these cultures for maintaining a historical record were only collected pre-transfer every 100 days to avoid potential disruption of ecological structure.

### DNA isolation and high-throughput sequencing

To track the mutational dynamics of the evolving populations, we collected 1 mL of each culture at days 90, 200, 300, 400, 600, 700, 800, and 900 of experimental evolution. DNA was extracted with the DNeasy UltraClean Microbial Kit (Qiagen 12224; formerly MO BIO UltraClean Microbial DNA Kit) before transferring the DNA samples to either The Hubbard Center for Genomic Analysis (University of New Hampshire), the Center for Genomics and Bioinformatics (Indiana University Bloomington), or the CLAS Genomics Facility (Arizona State University) for DNA library preparation and sequencing. Libraries were constructed by following a previous protocol that optimizes reagent usage for microbial genomes [85] with the Nextera DNA Library Preparation Kit (Illumina, FC-121-1030) and sequenced on an Illumina HiSeq 2500 (UNH) or an Illumina NextSeq 500 (Indiana; ASU). All sequencing produced paired-end data with a target depth of 100x coverage.

### Sequencing analysis

Quality control and mapping of NGS data were performed using previously described procedures [50]. Briefly, we used Cutadapt v.1.9.1 [86] to remove residual adapters and trim low-quality sequences, and resulting QCed sequencing reads were mapped to the *Escherichia coli* K-12 substr. MG1655 reference genome (NC_000913.3). Mutations, including SNPs, indels, IS-element insertions, and large deletions, as well as their frequencies were identified using Breseq v.0.30.2 with the predict-polymorphisms parameter setting [87]. Across all 128 genomic profiles (16 populations x 8 timepoints), we discarded eight low-quality genomic profiles resulting in 120 remaining profiles for analysis (**Table S1**).

For each experimental population at each sequenced time point, we estimated the level of genomic divergence by summing the observed frequencies of detected mutations. Genomic evolution rates were determined by the slope of the linear regression “genomic divergence ∼ generation numbers + 0” [53] using the function “lm” in R. the function “lm” in R. To determine the presence of co-existing subpopulations, we applied a clade-aware hidden Markov model (caHMM) using a modified version of previously released code [51] to the mutational dataset for each population [88].

### Gene-level analysis

Quantification of gene-level parallelism in evolved MMR- or WT populations was conducted as previously described [88]. Here, we identified a focal set of nonsynonymous mutations which included the fixed mutations from hidden Markov chain analysis [51] and the mutations showing DAF > 0.5 in at least two timepoints. Using this focal set of mutations, we calculated the *G*-score for each gene and the sum among all genes [53], where a larger sum suggests a higher gene-level parallelism. As the number of mutations in each focal set affects the expected distribution of the sum of *G*-scores, we performed 20,000 simulations to generate a null distribution for the sum of *G*-scores. The level of significance of the sum of *G*-scores was then evaluated by the *z* score, (the observed sum - the mean of simulated sums) / (the standard deviation of simulated sums). We also evaluated each gene to identify individual genes that were overrepresented for mutations by calculating the *P*-value for each gene as the proportion of 20,000 simulated *G*-scores larger or equal to the observed *G*-score. After Bonferroni correction, only the genes with corrected *P*-value < 0.05 are considered to be significantly overrepresented for nonsynonymous mutations. A list of genes significantly overrepresented for structural or nonsense mutations was acquired by a similar method focusing on indels, IS-element insertions, and nonsense mutations. As an additional method for assessing mutational parallelism, we also calculated the mean Bray-Curtis similarity across all pairs of experimental populations [54]. We generated a null distribution by performing 1,000 simulations that randomly sampled the nonsynonymous sites. The significance level was evaluated by a similar method as applied in the above *G*-score analysis, where *z-*scores were generated using the observed and simulated results.

To identify the presence of mutational order, for each population, we recorded the first sequencing timepoint at which a fixed mutation is detected in each gene that was determined to be overrepresented for non-synonymous and/or structural mutations. We then applied a linear model to assess if the time at which mutations first arise to detectable levels in different overrepresented genes was temporally restricted (timepoint ∼ gene) and assessed the statistical significance using ANOVA. We repeated this analysis, first within functional subsets of the data (i.e., regulation-, membrane-, or transport-associated genes) to determine how mutational order differed between gene functions and again with mutation types (timepoint ∼ mutation types) to determine if different mutation types were more common early or later in experimental evolution.

### Phenotyping of biofilm and motility

The ability to produce biofilm was assessed using a previously described microtiter plate biofilm assay [50, 56]. Briefly, 15 μL of overnight culture was inoculated into a single well of a non-tissue-treated 96-well plate (351172; Corning/Falcon) containing 150 μL of LB broth. Each population or engineered deletion mutant was measured for a total of eight replicates. As a control, each 96-well plate contained two columns (16 wells) of the *ΔaraBAD* WT ancestor and at least two columns (16 wells) of blanks. Biofilm was grown by incubating the 96-well plates without shaking for 24 h at 37°C. Following incubation, plates are thrice-washed with 1× phosphate-buffered saline (PBS) before staining with 0.1% crystal violet for 10 min (200 μL per well). After staining, plates are again thrice-washed with 1× PBS, inverted on an angle, and dried overnight. To quantify biofilm, we solubilized the biofilm-bound crystal violet with 30% acetate for 15 min (200 μL per well). Dissolved dye is then transferred to a fresh 96-well plate and absorbance is quantified with a Synergy H2 microplate spectrophotometer (Biotek) at 550 nm. Each assessed 96-well plate is treated as a batch. For each batch, adjusted biofilm values (x̂_Sample_) are calculated by subtracting the median absorbance value for the blank wells (x́_Blank_) from the individual absorbance values for each inoculated well (x_Sample_). Any negative or zero values produced after adjustment are replaced with 0.001 to allow for further normalization and statistical consideration. Relative biofilm values are then normalized to the WT ancestor by dividing each adjusted biofilm value (x̂_Sample_) by the median adjusted biofilm value of the WT ancestor (x́_WT,adj_) for that batch.

Motility of evolved *E. coli* populations and engineered deletion mutants was assessed by measuring growth on 0.35% motility plates [57]. *E. coli* samples were revived from −80°C frozen storage by overnight culture at 37°C in 15uL of LB. We dotted 1.5uL of revived samples on the surface of motility agar (5g tryptone, 1.75g granulated agar, 2.5g NaCl, 500 mL H_2_O - let dry for 24h). The dotted sample was left to dry on the surface of the motility agar for 30 minutes to allow the sample to set before the plates were transported to the incubator. Plates were incubated upright in a static 35°C incubator for 24h. After incubation, plates were imaged with an in-frame 20mm scale; initial spot area and motility area were measured with ImageJ (NIH). Relative motility was determined by calculating the difference of the radius of motile growth and the radius of the initial spot area. All samples were normalized by the mean motility of the WT ancestor for each batch.

### Construction of genetic mutants

Gene deletion strains were generated via P1 transduction by moving a kanamycin resistance cassette flanked by FRT sites from Keio collection single-gene knockout strains to the ancestral strain, PFM2 [89, 90]. For the single gene knockout strains, the *ΔfimE* strain was produced by transducing the Δ*fimE*781::kan allele into PFM2 from JW4276, the *ΔpaaX* strain was produced by transducing the Δ*paaX*783::kan allele into PFM2 from JW1394, the *ΔydcI* strain was produced by transducing the Δ*ydcI*783::kan allele into PFM2 from JW5226, and the *ΔhdfR* strain was produced by transducing the Δ*hdfR*785::kan allele into PFM2 from JW5607. In order to generate the *ΔhdfR/ΔfimE* double mutant strain, the kanamycin cassette was first excised from the *ΔhdfR* knockout strain [90]. Then, the Δ*fimE*781::kan allele was transduced from JW4276 into the aforementioned *ΔhdfR* knockout strain, generating the double mutant. Each transduction reaction was verified via PCR using primers that flanked each gene deletion site, followed by Sanger sequencing (**Table S5**).

### Fitness assessments for engineered deletion mutants

Pairwise competition assays were used to assess the relative fitness of engineered deletion mutants after 1-, 4- and 10-days of growth without resource replenishment, using a previously described method in which a different independent culture is assessed for each time point to preserve the spatial ecology of the co-culture and more accurately reflect the evolved environment [91]. As the culture experiences a large population reduction during death phase, we calculated the selection rate as a proxy for relative fitness with all selection rate calculations made relative to the WT PFM2 progenitor strain. Competitions were initiated by adding 50 µl of overnight culture for each competing strain to 10 ml of LB broth aiming for an approximate 50:50 starting proportion of each strain. Culture density was assessed for each overnight culture prior to setting up each competition by measuring the culture’s absorbance (600 nm). If large discrepancies between co-culture densities were noted, we adjusted their volumes to normalize for cell density. Immediately after preparing the competition co-culture, culture tubes were thoroughly vortexed before collecting a 100 µl aliquot for CFU counting and assessing the starting frequencies of each strain. Co-cultures were then incubated at 37 °C with 180 rpm shaking for the duration of the 1-, 4-, or 10-day competition after which the cultures vortexed and a 100 µl aliquot was collected to determine the final frequencies of each strain via CFU count.

To perform CFU counting, 100 µl aliquots of co-cultures were serially diluted in 1x PBS buffer and plated on TA agar (10 g/l tryptone, 1 g/l yeast extract, 5 g/l NaCl, 16 g/l agar, 10 g/l L-arabinose, 0.005% tetrazolium chloride). In order to distinguish between competing strains, it was ensured that one competitor in each co-culture had a neutral deletion of the Δ*araBAD* operon which produces dark red colonies on TA agar. Selection rate (r) was calculated as [ln(N_Mutant final_ - N_Mutant inital_)/(t)] - [ln(N_Ancestor final_ - N_Ancestor inital_)/(t)], the difference of the strain’s Malthusian parameters in co-culture [92], where N_final_ and N_initial_ are the final and initial CFU counts and t is the length of the competition in days.

### Data Availability

All code necessary to repeat genomic sequencing analysis and reproduce the figures can be found at https://github.com/BehringerLab/100-day-feast-famine, and all genomic sequencing reads from this study can be downloaded from SRA at NCBI (BioProject number PRJNA532905).

## Acknowledgements

We thank Gwyneth F. Boyer, Patricia Foster, Blane J. Hollingsworth, James B. McKinlay, Angelica Urquidez for their helpful comments and assistance. High-performance computing resources were provided and maintained by the National Center for Genome Analysis Support at Indiana University. This work was supported by Army Research Office grants W911NF-14-1-0411 (M.L.) and W911NF-21-1-0161 (M.G.B.) and National Institutes of Health grants F32GM123703 (M.G.B.) and R35GM122566 (M.L.).

## Supplemental Methods

### Backup population in experimental evolution

To avoid extinctions in the 100-day feast/famine conditions, populations were maintained in triplicate (1 main culture and 2 backup cultures) with only the main culture being used to seed new triplicate cultures every 100 days. In the case of a whole population extinction, one backup culture would be used to replace the main culture and the experiment would continue. This practice was only necessary in the first 200 days; after that time point, no additional whole-population extinction events occurred.

### Estimated generation numbers

The estimation of generation numbers in 100-day populations is based on the following two observations. First, pre-transfer population density in the culture tube was measured at day 200, 300, 400, 600, 700, 800, 900, by serial dilution in phosphate-buffered saline (PBS) before plating on LB agar for CFU counts and revealed that the population density does not significantly change throughout experimental evolution (**Fig. S6**). This suggests the carrying capacity of experimental populations is recovered within a resource-replenishment cycle after 1:10 dilution and the populations must have experienced at least 3.3 (log_2_10) generations between resource-replenishment cycles. Therefore, the first estimate of guaranteed generations (*g*_1_) per day equals 0.033. Second, we found that between resource-replenishment cycles novel mutations arise and are able to be fixed within one sequencing interval. Therefore, given the carrying capacity *K*, the population should at minimum experience log_2_*K* cell divisions in 100 days. Using estimated *K* = 3.71 x 10^7^ (**Fig. S6**), the second estimate of guaranteed generations (*g*_2_) per day equals 0.25. The final number of guaranteed generations (*g*) is then determined as *g*_2_, the larger value between *g*_1_ and *g*_2_.

---

**Supplemental Figure 1.**
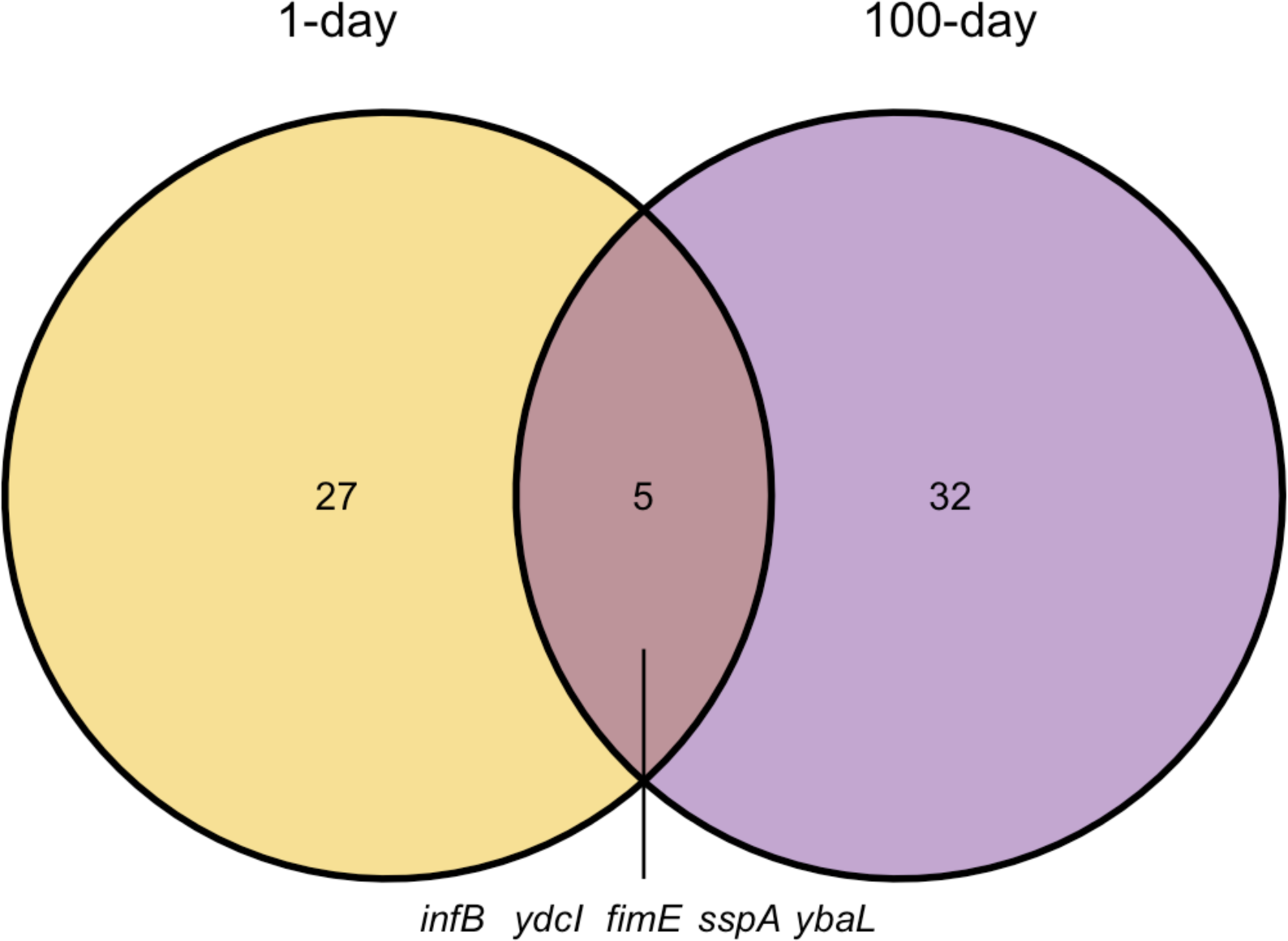
Venn diagram of genes overrepresented for mutations in 1- and 100-day feast/famine conditions. Overlap of genes determined to be overrepresented for mutations in populations evolving to 1-day (yellow/ left circle; Behringer and Ho, et al. 2022 [50]) and 100-day (purple/ right circle; Behringer and Ho, et al. this study). Numbers represent the number of genes in each group, with genes overrepresented for mutations in both treatments labelled in the intersection. Circles are drawn as evenly sized and not drawn to scale with number of genes.

**Supplemental Figure 2.**
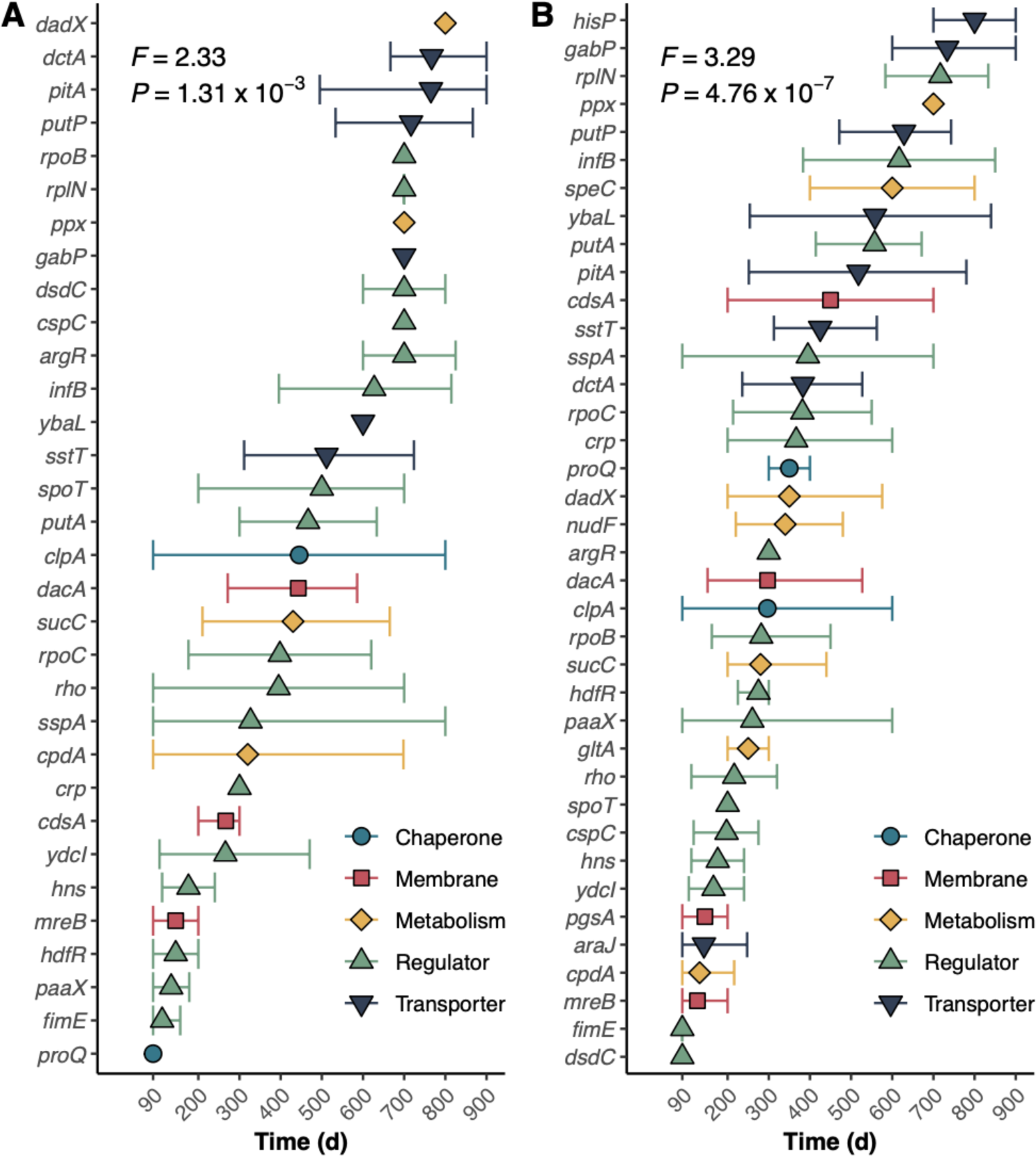
Mutational parallelism is significant in (A) WT and (B) MMR-populations. Colors and filled shapes indicate the annotated function of each gene. For both plots, filled shapes indicate the mean timepoint at which the mutation is first detected in a population. Bars represent the 95% confidence interval of the mean.

**Supplemental Figure 3.**
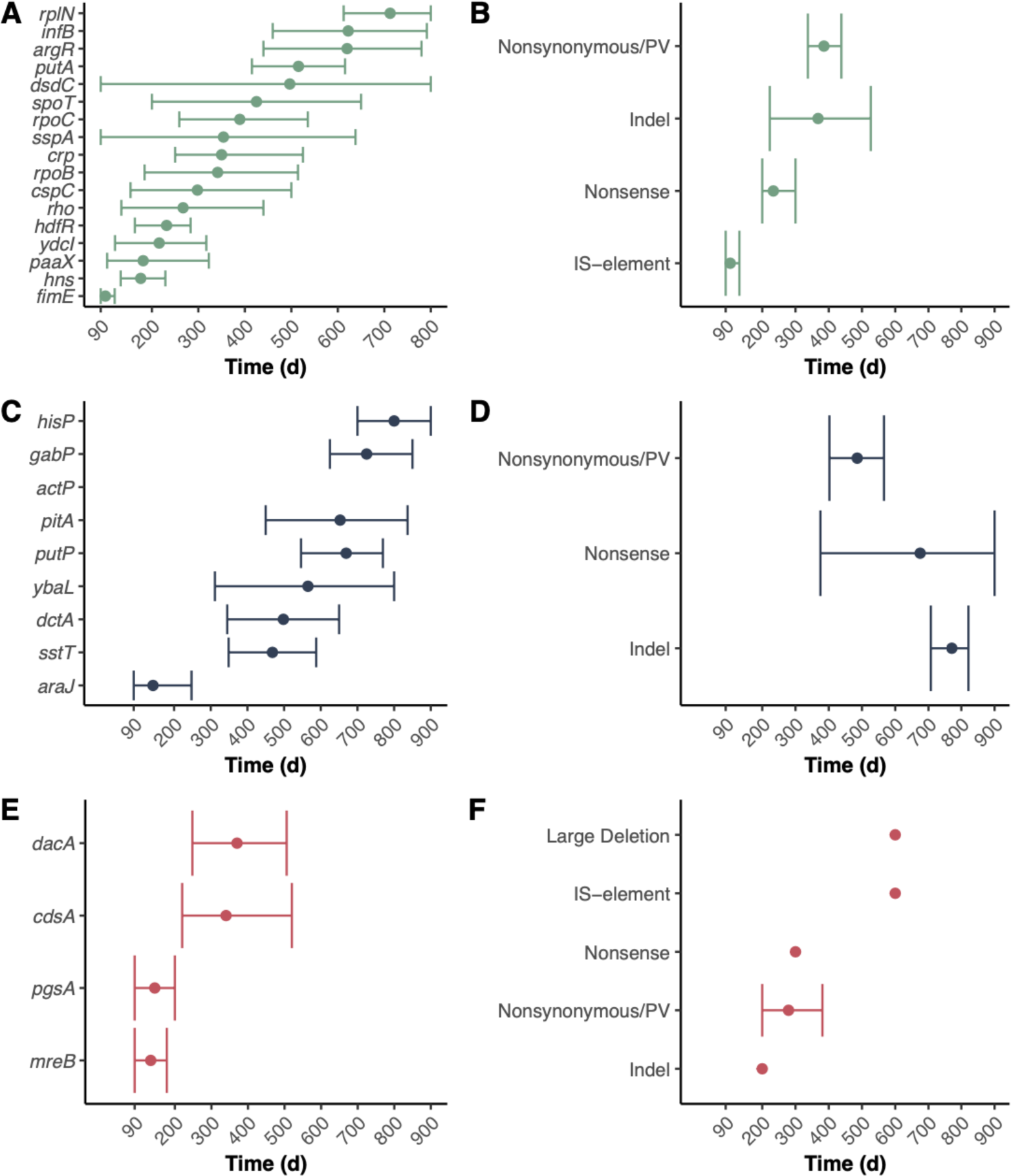
Mutational parallelism by gene type. Plots are separated by genes annotated as regulators (A, B; green), genes annotated as transporters (C, D; navy), and genes annotated as associated with the cell envelope (E, F, red). For all plots, filled circles indicate the mean timepoint at which the mutation is first detected in a population. Bars represent the 95% confidence interval of the mean.

**Supplemental Figure 4.**
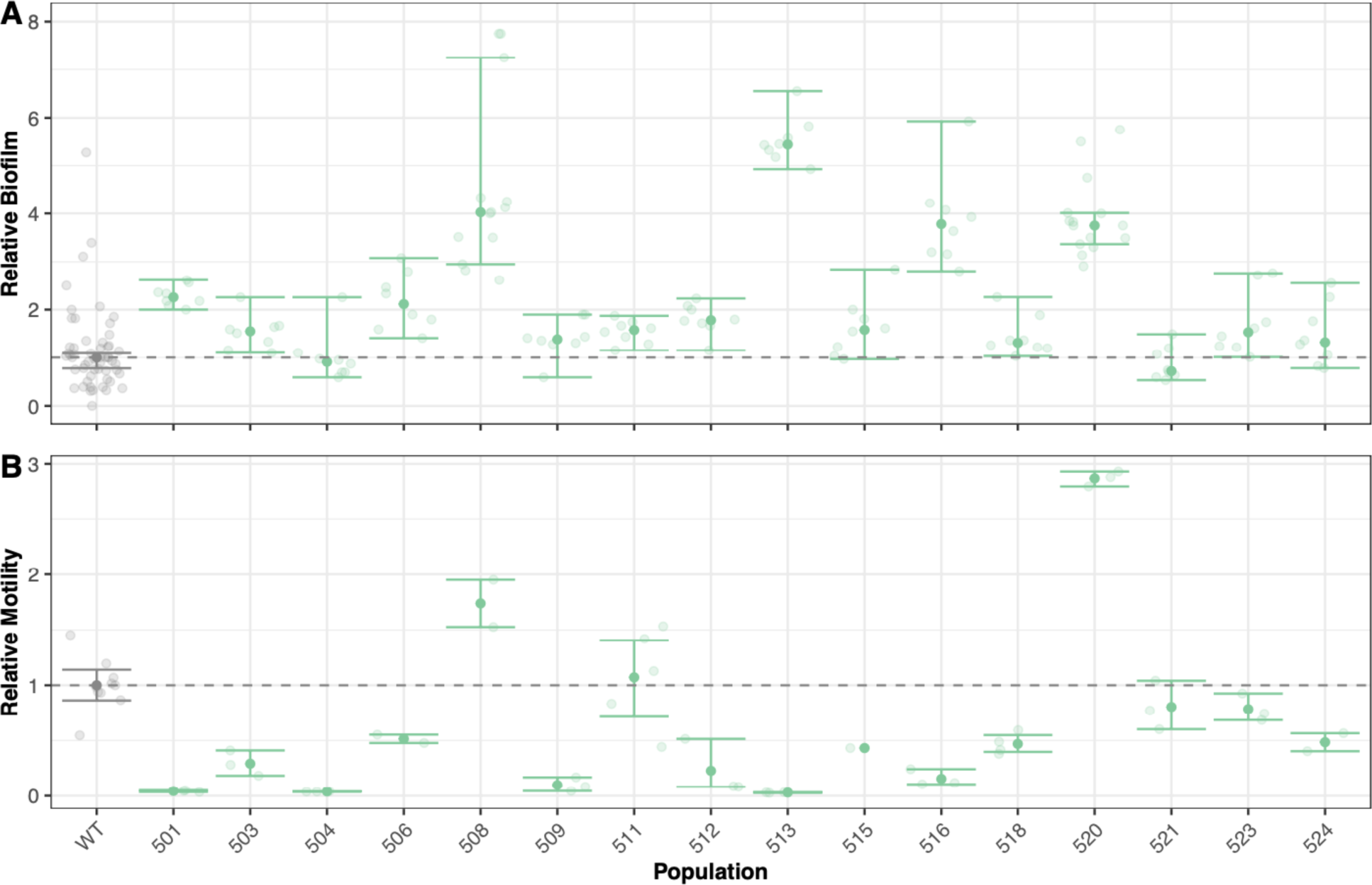
Measurements of biofilm (A) and motility (B) for all evolved populations. Transparent circles indicate the individual data points for each population (green). All relative measurements are normalized to the mean value of the WT ancestor (gray). For both plots, filled circles indicate the mean value for each population and error bars represent the 95% confidence interval of the mean.

**Supplemental Figure 5.**
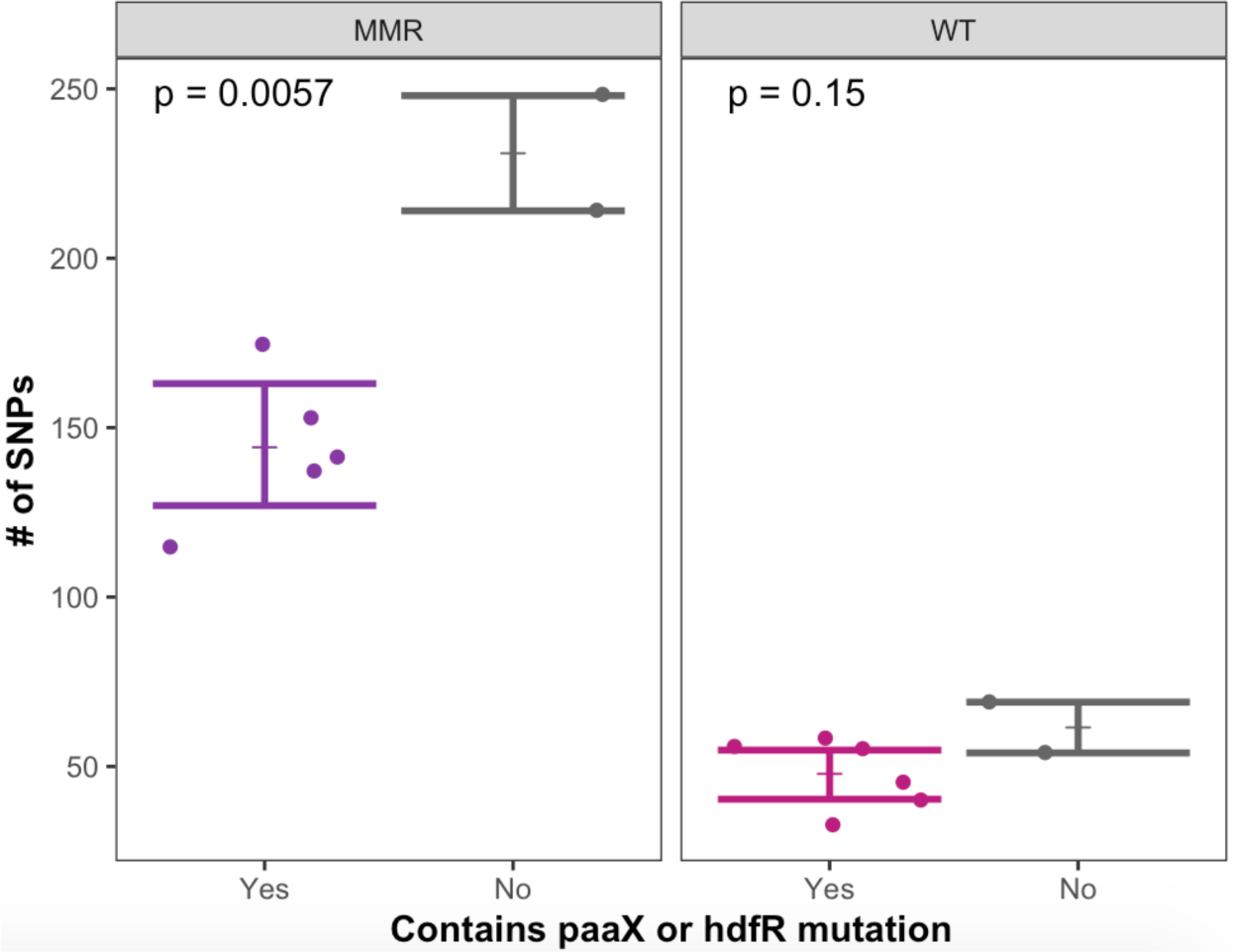
SNP Count varies based on presence of mutation in *paaX* or *hdfR*. Filled circles indicate the # of SNPs at a frequency of > 10% in each evolved population after 900-days. Horizontal bar represents the mean SNP count and error bars represent the 95% confidence interval of the mean. P-value is calculated from a Welch’s T-test.

**Supplemental Figure 6.**
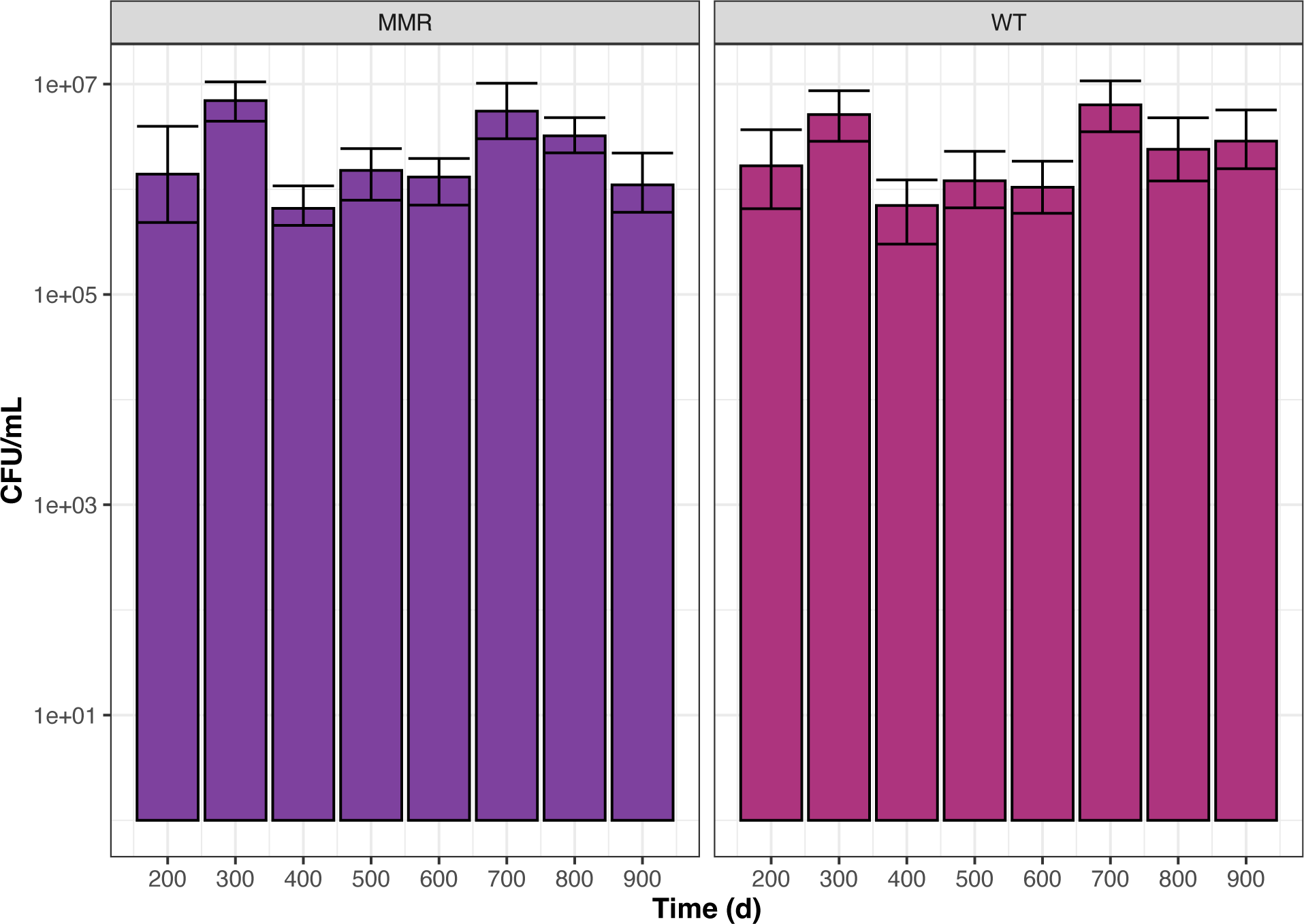
Population density of evolved populations. Bars represent mean pre-transfer population density of WT and MMR-populations for each transfer during experimental evolution. Error bars represent ± 95% confidence interval of the mean.

## Literature Cited

1. Hobbie JE, Hobbie EA. Microbes in nature are limited by carbon and energy: the starving-survival lifestyle in soil and consequences for estimating microbial rates. Front Microbiol 2013; 4: 324.

2. Zhang YJ, Rubin EJ. Feast or famine: the host-pathogen battle over amino acids: Host-pathogen battle over amino acids. Cell Microbiol 2013; 15: 1079–1087.

3. Reese AT, Pereira FC, Schintlmeister A, Berry D, Wagner M, Hale LP, et al. Microbial nitrogen limitation in the mammalian large intestine. Nat Microbiol 2018; 3: 1441–1450.

4. Johnson JR, Russo TA. Extraintestinal pathogenic *Escherichia coli*: ‘the other bad E coli’. J Lab Clin Med 2002; 139: 155–162.

5. Smith JL, Fratamico PM, Gunther NW. Extraintestinal pathogenic *Escherichia coli*. Foodborne Pathog Dis 2007; 4: 134–163.

6. Brown SP, Cornforth DM, Mideo N. Evolution of virulence in opportunistic pathogens: generalism, plasticity, and control. Trends Microbiol 2012; 20: 336–342.

7. Jang J, Hur H-G, Sadowsky MJ, Byappanahalli MN, Yan T, Ishii S. Environmental *Escherichia coli*: ecology and public health implications-a review. J Appl Microbiol 2017; 123: 570–581.

8. Barton IS, Fuqua C, Platt TG. Ecological and evolutionary dynamics of a model facultative pathogen: Agrobacterium and crown gall disease of plants. Environ Microbiol 2018; 20: 16–29.

9. Vasi FK, Lenski RE. Ecological strategies and fitness tradeoffs in *Escherichia coli* mutants adapted to prolonged starvation. J Genet 1999; 78: 43–49.

10. Ferenci T. Maintaining a healthy SPANC balance through regulatory and mutational adaptation. Mol Microbiol 2005; 57: 1–8.

11. Ying B-W, Honda T, Tsuru S, Seno S, Matsuda H, Kazuta Y, et al. Evolutionary consequence of a trade-off between growth and maintenance along with ribosomal damages. PLoS ONE 2015; 10: e0135639.

12. Ferenci T. Trade-off mechanisms shaping the diversity of bacteria. Trends Microbiol 2016; 24: 209–223.

13. Zimmer DP, Soupene E, Lee HL, Wendisch VF, Khodursky AB, Peter BJ, et al. Nitrogen regulatory protein C-controlled genes of *Escherichia coli* : Scavenging as a defense against nitrogen limitation. Proc Natl Acad Sci USA 2000; 97: 14674–14679.

14. Switzer A, Burchell L, McQuail J, Wigneshweraraj S. The adaptive response to long-term nitrogen starvation in *Escherichia coli* requires the breakdown of allantoin. J Bacteriol 2020; 202: e00172–20.

15. Taddei F, Matic I, Radman M. cAMP-dependent SOS induction and mutagenesis in resting bacterial populations. Proc Natl Acad Sci USA 1995; 92: 11736–11740.

16. Loewe L, Textor V, Scherer S. High deleterious genomic mutation rate in stationary phase of *Escherichia coli*. Science 2003; 302: 1558–60.

17. Sniegowski P. Evolution: bacterial mutation in stationary phase. Curr Biol 2004; 14: R245–6.

18. Foster PL. Stress-induced mutagenesis in bacteria. Critical Reviews in Biochemistry and Molecular Biology 2007; 42: 373–397.

19. Matic I. Stress-induced mutagenesis in bacteria. In: Mittelman D (ed). Stress-Induced Mutagenesis. 2013. Springer New York, New York, NY, pp 1–19.

20. Cohen D. Optimizing reproduction in a randomly varying environment. Journal of Theoretical Biology 1966; 12: 119–129.

21. Ellner S. ESS germination strategies in randomly varying environments. I. Logistic-type models. Theoretical Population Biology 1985; 28: 50–79.

22. Kaprelyants AS, Gottschal JC, Kell DB. Dormancy in non-sporulating bacteria. FEMS Microbiology Letters 1993; 104: 271–286.

23. Lennon JT, Jones SE. Microbial seed banks: the ecological and evolutionary implications of dormancy. Nat Rev Microbiol 2011; 9: 119–130.

24. Shoemaker WR, Lennon JT. Evolution with a seed bank: The population genetic consequences of microbial dormancy. Evol Appl 2018; 11: 60–75.

25. Piggot PJ, Hilbert DW. Sporulation of *Bacillus subtilis*. Current Opinion in Microbiology 2004; 7: 579–586.

26. Mutlu A, Kaspar C, Becker N, Bischofs IB. A spore quality–quantity tradeoff favors diverse sporulation strategies in Bacillus subtilis. ISME J 2020; 14: 2703–2714.

27. Rathmann I, Förster M, Yüksel M, Horst L, Petrungaro G, Bollenbach T, et al. Distribution of fitness effects of cross-species transformation reveals potential for fast adaptive evolution. ISME J 2023; 17: 130–139.

28. Browne HP, Almeida A, Kumar N, Vervier K, Adoum AT, Viciani E, et al. Host adaptation in gut Firmicutes is associated with sporulation loss and altered transmission cycle. Genome Biol 2021; 22: 204.

29. Ferenci T. Hungry bacteria - definition and properties of a nutritional state. Environ Microbiol 2001; 3: 605–611.

30. Navarro Llorens JM, Tormo A, Martínez-García E. Stationary phase in gram-negative bacteria. FEMS Microbiol Rev 2010; 34: 476–495.

31. Yan J, Nadell CD, Bassler BL. Environmental fluctuation governs selection for plasticity in biofilm production. ISME J 2017; 11: 1569–1577.

32. Valencia EY, Barros JP, Ferenci T, Spira B. A broad continuum of E. coli traits in nature associated with the trade-off between self-preservation and nutritional competence. Microb Ecol 2022; 83: 68– 82.

33. Van Elsas JD, Semenov AV, Costa R, Trevors JT. Survival of *Escherichia coli* in the environment: fundamental and public health aspects. The ISME journal 2011; 5: 173–183.

34. Flint KP. The long-term survival of *Escherichia coli* in river water. Journal of Applied Bacteriology 1987; 63: 261–270.

35. Brennan FP, O’Flaherty V, Kramers G, Grant J, Richards KG. Long-term persistence and leaching of *Escherichia coli* in temperate maritime soils. Appl Environ Microb 2010; 76: 1449–1455.

36. Zambrano MM, Siegele DA, Almirón M, Tormo A, Kolter R. Microbial competition: *Escherichia coli* mutants that take over stationary phase cultures. Science 1993; 259: 1757–1760.

37. Finkel SE. Long-term survival during stationary phase: evolution and the GASP phenotype. Nat Rev Microbiol 2006; 4: 113–120.

38. Avrani S, Bolotin E, Katz S, Hershberg R. Rapid genetic adaptation during the first four months of survival under resource exhaustion. Mol Biol Evol 2017; 34: 1758–1769.

39. Ratib NR, Seidl F, Ehrenreich IM, Finkel SE. Evolution in long-term stationary-phase batch culture: emergence of divergent *Escherichia coli* lineages over 1,200 days. mBio 2021; 12: e03337–20.

40. Vasi F, Travisano M, Lenski RE. Long-Term experimental evolution in *Escherichia coli*. II. Changes in life-history traits during adaptation to a seasonal environment. The American Naturalist 1994; 144: 432–456.

41. Kram KE, Geiger C, Ismail WM, Lee H, Tang H, Foster PL, et al. Adaptation of *Escherichia coli* to long-term serial passage in complex medium: evidence of parallel evolution. mSystems 2017; 2: e00192–16.

42. Behringer MG, Choi BI, Miller SF, Doak TG, Karty JA, Guo W, et al. *Escherichia coli* cultures maintain stable subpopulation structure during long-term evolution. Proc Natl Acad Sci USA 2018; 115.

43. Farrell MJ, Finkel SE. The growth advantage in stationary-phase phenotype conferred by rpoS mutations is dependent on the pH and nutrient environment. J Bacteriol 2003; 185: 7044–7052.

44. Zinser ER, Kolter R. Prolonged stationary-phase incubation selects for lrp mutations in *Escherichia coli* K-12. J Bacteriol 2000; 182: 4361–5.

45. Zambrano MM, Kolter R. GASPing for life in stationary phase. Cell 1996; 86: 181–4.

46. Finkel SE, Kolter R. Evolution of microbial diversity during prolonged starvation. Proc Natl Acad Sci U S A 1999; 96: 4023–7.

47. Chib S, Ali F, Seshasayee ASN. Genomewide mutational diversity in *Escherichia coli* population evolving in prolonged stationary phase. Msphere 2017; 2.

48. Lee H, Popodi E, Tang HX, Foster PL. Rate and molecular spectrum of spontaneous mutations in the bacterium *Escherichia coli* as determined by whole-genome sequencing. Proc Natl Acad Sci U S A 2012; 109: E2774–E2783.

49. Wei W, Ho W-C, Behringer MG, Miller SF, Bcharah G, Lynch M. Rapid evolution of mutation rate and spectrum in response to environmental and population-genetic challenges. Nat Commun 2022; 13: 4752.

50. Behringer MG, Ho W-C, Meraz JC, Miller SF, Boyer GF, Stone CJ, et al. Complex ecotype dynamics evolve in response to fluctuating resources. mBio 2022; 13: e03467–21.

51. Good BH, McDonald MJ, Barrick JE, Lenski RE, Desai MM. The dynamics of molecular evolution over 60,000 generations. Nature 2017; 551: 45–50.

52. Press MO, Hall AN, Morton EA, Queitsch C. Substitutions are boring: some arguments about parallel mutations and high mutation rates. Trends in Genetics 2019; 35: 253–264.

53. Tenaillon O, Barrick JE, Ribeck N, Deatherage DE, Blanchard JL, Dasgupta A, et al. Tempo and mode of genome evolution in a 50,000-generation experiment. Nature 2016; 536: 165–70.

54. Turner CB, Marshall CW, Cooper VS. Parallel genetic adaptation across environments differing in mode of growth or resource availability. Evolution letters 2018; 2: 355–367.

55. Gillespie JH. Genetic drift in an infinite population: The pseudohitchhiking model. Genetics 2000; 155: 909–19.

56. O’Toole GA. Microtiter dish biofilm formation assay. JoVE 2011; 2437.

57. Wolfe AJ, Berg HC. Migration of bacteria in semisolid agar. Proc Natl Acad Sci USA 1989; 86: 6973–6977.

58. Nadell CD, Bassler BL. A fitness trade-off between local competition and dispersal in *Vibrio cholerae* biofilms. Proc Natl Acad Sci USA 2011; 108: 14181–14185.

59. Penterman J, Nguyen D, Anderson E, Staudinger BJ, Greenberg EP, Lam JS, et al. Rapid evolution of culture-impaired bacteria during adaptation to biofilm growth. Cell Reports 2014; 6: 293–300.

60. Lowery NV, McNally L, Ratcliff WC, Brown SP. Division of labor, bet hedging, and the evolution of mixed biofilm investment strategies. mBio 2017; 8: e00672–17.

61. van Ditmarsch D, Boyle KE, Sakhtah H, Oyler JE, Nadell CD, Déziel É, et al. Convergent evolution of hyperswarming leads to impaired biofilm formation in pathogenic bacteria. Cell Reports 2013; 4: 697–708.

62. Nyström T. Growth versus maintenance: a trade-off dictated by RNA polymerase availability and sigma factor competition?: Transcriptional trade-offs. Molecular Microbiology 2004; 54: 855–862.

63. Biselli E, Schink SJ, Gerland U. Slower growth of *Escherichia coli* leads to longer survival in carbon starvation due to a decrease in the maintenance rate. Molecular Systems Biology 2020; 16: e9478.

64. Abram F, Arcari T, Guerreiro D, O’Byrne CP. Evolutionary trade-offs between growth and survival: The delicate balance between reproductive success and longevity in bacteria. Adv Microb Physiol 2021; 79: 133–162.

65. Ercan O, Den Besten HMW, Smid EJ, Kleerebezem M. The growth-survival trade-off is hard-wired in the *Lactococcus lactis* gene regulation network. Environ Microbiol Rep 2022; 14: 632–636.

66. Orgogozo V. Replaying the tape of life in the twenty-first century. Interface Focus 2015; 5: 20150057.

67. Woods R, Schneider D, Winkworth CL, Riley MA, Lenski RE. Tests of parallel molecular evolution in a long-term experiment with *Escherichia coli*. Proc Natl Acad Sci USA 2006; 103: 9107–9112.

68. Deatherage DE, Kepner JL, Bennett AF, Lenski RE, Barrick JE. Specificity of genome evolution in experimental populations of *Escherichia coli* evolved at different temperatures. Proc Natl Acad Sci USA 2017; 114.

69. Santos-Lopez A, Marshall CW, Scribner MR, Snyder DJ, Cooper VS. Evolutionary pathways to antibiotic resistance are dependent upon environmental structure and bacterial lifestyle. eLife 2019; 8: e47612.

70. Dragosits M, Mozhayskiy V, Quinones-Soto S, Park J, Tagkopoulos I. Evolutionary potential, cross-stress behavior and the genetic basis of acquired stress resistance in *Escherichia coli*. Molecular Systems Biology 2013; 9: 643.

71. Mani GS, Clarke BC. Mutational order: a major stochastic process in evolution. Proc R Soc Lond B 1990; 240: 29–37.

72. Schluter D. Evidence for ecological speciation and its alternative. Science 2009; 323: 737–741.

73. Kent DG, Green AR. Order matters: the order of somatic mutations influences cancer evolution. Cold Spring Harb Perspect Med 2017; 7: a027060.

74. Huang B-H, Lin Y-C, Huang C-W, Lu H-P, Luo M-X, Liao P-C. Differential genetic responses to the stress revealed the mutation-order adaptive divergence between two sympatric ginger species. BMC Genomics 2018; 19: 692.

75. Cooper TF. Empirical insights into adaptive landscapes from bacterial experimental evolution. In: Svensson E, Calsbeek R (eds). The Adaptive Landscape in Evolutionary Biology. 2013. Oxford University Press, p 0.

76. Ono J, Gerstein AC, Otto SP. Widespread genetic incompatibilities between first-step mutations during parallel adaptation of *Saccharomyces cerevisiae* to a common environment. PLoS Biol 2017; 15: e1002591.

77. Harmand N, Gallet R, Martin G, Lenormand T. Evolution of bacteria specialization along an antibiotic dose gradient. Evolution Letters 2018; 2: 221–232.

78. Oxman E, Alon U, Dekel E. Defined order of evolutionary adaptations: experimental evidence. Evolution 2008; 62: 1547–1554.

79. Buswell CM, Herlihy YM, Lawrence LM, McGuiggan JTM, Marsh PD, Keevil CW, et al. Extended survival and persistence of *Campylobacter* spp. in water and aquatic biofilms and their detection by immunofluorescent-antibody and -rRNA staining. Appl Environ Microbiol 1998; 64: 733–741.

80. Switzer A, Evangelopoulos D, Figueira R, De Carvalho LPS, Brown DR, Wigneshweraraj S. A novel regulatory factor affecting the transcription of methionine biosynthesis genes in *Escherichia coli* experiencing sustained nitrogen starvation. Microbiology 2018; 164: 1457–1470.

81. Reitzer L. Biosynthesis of glutamate, aspartate, asparagine, L -alanine, and D -alanine. EcoSal Plus 2004; 1: ecosalplus.3.6.1.3.

82. Al Mamun AAM, Lombardo M-J, Shee C, Lisewski AM, Gonzalez C, Lin D, et al. Identity and function of a large gene network underlying mutagenic repair of DNA breaks. Science 2012; 338: 1344–1348.

83. Yang H, Wolff E, Kim M, Diep A, Miller JH. Identification of mutator genes and mutational pathways in *Escherichia coli* using a multicopy cloning approach: Discover mutator genes by multicopy cloning. Molecular Microbiology 2004; 53: 283–295.

84. Tenaillon O, Denamur E, Matic I. Evolutionary significance of stress-induced mutagenesis in bacteria. Trends in Microbiology 2004; 12: 264–270.

85. Baym M, Kryazhimskiy S, Lieberman TD, Chung H, Desai MM, Kishony R. Inexpensive multiplexed library preparation for megabase-sized genomes. PLoS One 2015; 10: e0128036.

86. Martin M. Cutadapt removes adapter sequences from high-throughput sequencing reads. EMBnet.journal 2011; 17: 10–12.

87. Deatherage DE, Barrick JE. Identification of mutations in laboratory-evolved microbes from next-generation sequencing data using breseq. Methods Mol Biol 2014; 1151: 165–88.

88. Ho W-C, Behringer MG, Miller SF, Gonzales J, Nguyen A, Allahwerdy M, et al. Evolutionary dynamics of asexual hypermutators adapting to a novel environment. Genome Biology and Evolution 2021; 13: evab257.

89. Baba T, Ara T, Hasegawa M, Takai Y, Okumura Y, Baba M, et al. Construction of *Escherichia coli* K-12 in-frame, single-gene knockout mutants: the Keio collection. Molecular Systems Biology 2006; 2: 2006.0008.

90. Saragliadis A, Trunk T, Leo JC. Producing gene deletions in *Escherichia coli* by P1 transduction with excisable antibiotic resistance cassettes. JoVE 2018; 58267.

91. Worthan SB, McCarthy RDP, Behringer MG. Case studies in the assessment of microbial fitness: seemingly subtle changes can have major effects on phenotypic outcomes. J Mol Evol 2023; 91: 311–324.

92. Lenski RE, Rose MR, Simpson SC, Tadler SC. Long-term experimental evolution in *Escherichia coli*. I. Adaptation and divergence during 2,000 generations. The American Naturalist 1991; 138: 1315–1341.

